# Surveillance of airborne plant disease dissemination at continental scale using air mass trajectory analysis and network theory

**DOI:** 10.1101/2021.06.04.447025

**Authors:** Andrea Radici, Davide Martinetti, Daniele Bevacqua

## Abstract

Sustainable management of plant disease outbreaks in agriculture is one of the main challenges of the next years to restore economic and environmental viability of farming practices. Improving early-detection capabilities and disease surveillance is increasingly seen as an obligate step to design appropriate and effective prophylactic measures. In this context, plant diseases caused by wind-dispersed pathogens represent an interesting case of study, since they are particularly complex and hard to observe directly, especially if compared to other dissemination means, and demand for a multidisciplinary approach to be dealt with. Wind dispersal could imply a geographic differentiation in pathogens spreading potential, due to the emerging of local meteorological features. In this work we analyze the spatio-temporal patterns of wind connectivity in Europe and the Mediterranean basin in order to identify possible pathways of *Puccinia graminis* spores, the causal agent of stem rust of wheat. By running backwards Lagrangian simulations merging a biological layer coupled with a pathogen viability model, we investigate possible long-distance connections between regions in the study area across different seasons. We characterized these regions in terms of network centrality indicators to identify possible spreaders of stem rust of wheat, founding that Central and Western European regions appears to provide highest connectivity for the spread of *P. graminis*.

## I. Introduction

Plant disease outbreaks in the agricultural context are a major sustainability concern because of two main reasons. The first and most obvious is directly related to food security, since outbreaks can cause harvest and subsequent economic losses to farmers due to damaged crops. This threat has been a constant trough out human history [1]. Nevertheless, in recent decades a second reason of concern has emerged, related to the environmental and social sustainability of farming practices. The widespread use of phytosanitary products, in the attempt of ensuring short and middle time crop production, may generate negative spillover effects affecting biodiversity, environmental pollution and health-related issues [2]. Ensuring the economic, social and environmental viability of farming practices is widely recognised as one of the main challenges agriculture has to face in the next years [3], [4].

In the context of the epidemic surveillance of plants diseases, wind-dispersed fungal pathogens have emerged as an interesting case of study. Long-distance aerial transport of pathogens can be responsible for different outcomes. In may lead to the first introduction of a pathogen in a region or continent, or to the seasonal reintroduction of a pathogen that cannot survive over the nongrowing season and, finally, to the spread of the disease between farms [5]. The reason behind this interest lies in the possibility of approaching the dispersal mechanisms by means of multiple disciplines. In recent times, aerobiology of spores and pollen has been studied by means of Lagrangian approaches, such as the Hybrid Single-Particle Lagrangian Integrated Trajectory model (HYSPLIT) [7], [8], in order to reconstruct episodes causing the introduction of pathogens in a new geographical context [6] or trying to explain anomalous concentrations of biological particles [9]. Few works tried to link possible vulnerabilities of regions in terms of epidemic outbreaks with respect to wind patterns [10]–[12].

The purpose of this work is to identify geographical hotspots for the spread of *Puccinia graminis* spores in the European and Mediterranean context due to wind dissemination. *P. graminis* is the fungal causal agent of stem rust of wheat (also known as stem rust) that is mainly dispersed by wind [13], [19]. In spite of the operations of containment and removal of obliged hosts carried out in the past decades, new varieties of stem rust of wheat have recently emerged, the TTKSK (Ug99) variety in Uganda being the most famous case [14], [15]. We first identified the main biotic and abiotic factors characterising *P. graminis* dispersal and settlement, and used such information to run backwards Lagrangian trajectory simulations to identify suitable connections between release locations and deposition sites. We then used the formalism of complex connectivity networks to support the identification of possible spreaders by means of centrality indicators.

## II. Materials and Methods

### A. Description of the case study and geographic domain

In order to study wind connectivity in the European and Mediterranean context, a first domain, spanning from the northern shores of Africa to the Scandinavian Peninsula, and from Iceand to the Ural mountains, was defined (27 °N, 71 °N; 26° W, 55° E). This domain was subsequently divided in a grid of 14,080 cells, measuring 0.5° × 0.5° (approximately 2,000 km^2^). This cell dimension was chosen since it corresponds to the resolution of meteorological data of the GDAS grid used by the HYSPLIT software for computing wind trajectories. This gridded domain was further filtered assuming a land use criteria: only cells covered with a surface of at least 1 % of possible susceptible host – i.e., wheat fields– were maintained in this analysis, thus excluding those regions where the host surface is less consistent. Land use raster provided by Earthstat database [16], referring to land covers in years 1997-2003, was used. Due to some missing data in specific locations, an alternative database has been used to identify wheat surface in Sicily (Italy) and Cyprus. Corine Land Cover surfaces of class 2.1.2 (i.e., “Non-irrigated arable land”) was used as a proxy of wheat surfaces [17]. The same filtering criterion (cells covered with a surface at least 1 % of class 2.1.2) was kept to select only cells covered with a consistent host surface. As a result, 3,870 cells were extracted from the original gridded domain, i.e. 27% of the initial domain (Fig. 1).

**Fig. 1.**
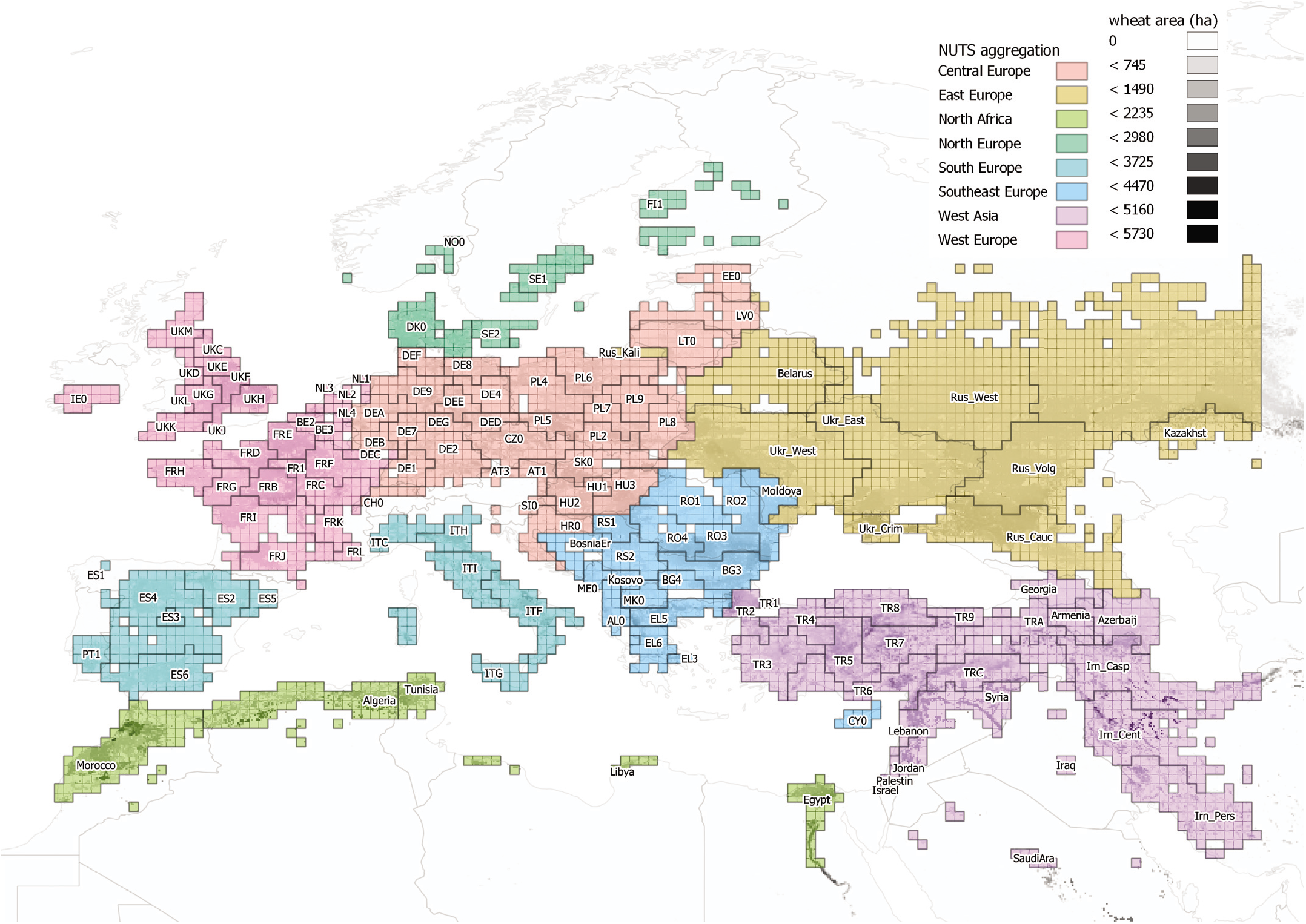
Domain cells aggregation into NUTS1 units, or equivalent, coloured by macroregions.

### B. Lagrangian simulations settings

Backward air-mass trajectories of 72 hours of duration were calculated using the HYSPLIT software from January 1, 2013 to December 31, 2018 (as in [11]). The centroids of the previously defined cells chosen across the study area were used as the arrival sites of the air-mass trajectories. Simulations were run every 3 h starting from midnight, so that 8 daily trajectories were obtained from each arrival site. The arrival simulation altitude was set to be 500 m above ground level [9], as an intermediate value to be below the clouds level and above the mostly mixed layers of the troposphere. The altitude of the centroids was calculated using a digital elevation model [18]. The total number of computed trajectories was 67,802,400. In addition to the three-dimensional spatial trajectory (longitude, latitude and altitude), other meteorological and atmospheric variables, such as rain, solar radiation, relative humidity, pressure, mixed layer depth were also recorder along all trajectories.

### C. Selection of suitable trajectories

In order to consider only those air-masses movements that could effectively lead to a spore dissemination event we imposed a set of filtering criteria inspired by the physical process of spore release, transport and deposition. This selection tried to account for the so called “disease triangle”, according to which the buildup of a pathogen requires the contemporary presence of (a) a susceptible host plant, (b) a virulent and viable pathogen and (c) favourable environmental conditions conducive to infection [20]. The filtering criteria were generally chosen between those suggested by the relevant literature to represent the dissemination phenomena of *P. graminis* spores:

1. **Wet deposition** - Fungal spores can be deposited via both dry and wet depositions mechanism. It has been suggested that during a given year, the contributions of wet and dry deposition are roughly equal in number, even though there are more dry hours than wet hours within an average season [21]. However, wet deposition may provide a better environment for the development of infection than dry deposition, since a certain level of soil and canopy humidity are needed for the spores to germinate [13]. Previous modelling works on wheat stem rust considered irrigation or a soil relative humidity higher than 90% as a necessary condition for infection [11]. As consequence, we consider only dissemination events that occur under wet deposition. This criterion has been traduced into filtering all those trajectories whose arrival sites satisfies the condition for which the precipitation intensity *r* (mm/h) *>* 0. All other trajectories whose arrival sites does not satisfy this condition were hence discarded.
2. **Spore survival to radiation exposure** - Fungal spores must endure exposure to harmful radiations, extremes temperatures and other harsh environmental conditions while airborne [13]. However, the limiting factor affecting *P. graminis* spore survival in the atmosphere is considered to be the exposure to solar radiation [11]. In this work we estimated the probability *P*(*SR*) of survival to harmful cumulative solar radiation *SR* (MJ) of a concentration of spores along a trajectory as:

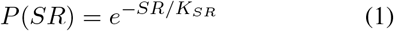 This equation has been calibrated in terms of the co-efficient *K*_*SR*_, representing the sensibility of spores to solar radiation for few species of fungi [20]. Since it has not been calibrated for *P. graminis*, a mean critical dose from other species was used, as already suggested by literature [6]. A value of *K*_*SR*_ = 14.0 MJ was adopted, and the probability of survival to solar radiation was calculated between each arrival site and the possible release locations using eq. 1. As a criterion to discard those unsuitable release locations characterised by a low probability of survival to harmful solar radiation, a boolean condition was adopted: only those locations whose *P*(*SR*) was larger than 0.1 were kept, and the remaining were discarded.
3. **Spore loss to rain washout** - A secondary phenomenon influencing the probability of dissemination of spores is the possible washout of the spores from the air mass due to precipitation events along the trajectory. In fact, rain droplets can scavenge particulate matter from the atmosphere to the ground, according to various parameters related to particles aerodynamic characteristics, while it also known that certain microorganisms can prompt the formation of ice nuclei in clouds conducive to rain [22]. Hence, a similar approach to the one followed with the solar radiation was adopted. Isard *et al*. [6] suggested an exponential relation describing the fraction of spores (here interpreted as a single spore survival probability *P*(*R*)), remaining in the atmosphere as a function of the cumulative rainfall *R* (mm) along the trajectory:

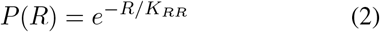 As in eq. 1, *K*_*R*_, represents the loss of spores to cumulative rainfall along the trajectory. A value of *K*_*SR*_ = 25.4 mm was adopted [6]. The same boolean condition applied for solar radiation was used for rainfall: possible release locations whose *P*(*R*) was larger than 0.1 were kept, while the others were discarded.
4. **Interception with the ground** - A criterion was set to discard those air masses whose altitude was considered to be too elevated for actively catch spores. It is known that spores are released at the canopy and ground level, but those progressively escape the lower layers of the troposphere and are dispersed in the air column within the range of few kilometers under turbulent atmospheric conditions [5], [13]. Since the aim of the study is to investigate long-distance dispersal at a large scale, we set another filtering criterion to identify suitable air-mass altitudes with respect to the height of the planetary boundary layer along the trajectory. Those locations crossed by an air mass trajectory within the mixed layer depth (MLD; therefore, by air masses able to drag spores distributed in the air column) were considered as suitable, while the others were discarded.
5. **Timing of spore release** - Spores are released by fungi according to several mechanism [5], [13], [23], [24]. It is known that *P. graminis* spores are passively released, which means that they leave the plant canopy mainly because of a mechanical stimulation (such as sudden strong wind gusts) which allows the spore to win the electrostatic bounding forces [24] and pass from the laminar layer close to the leaf surface into the turbulent layer within the crop [13]. In literature, the decrease of bounding forces in passively released spores has been correlated with a decrease in relative humidity and an increase of solar radiation [24], which usually occurs in the morning and in the early afternoon. Due to the difficulty in detecting sudden strong gusts from meteorological data (in opposition to the time-averaged wind reconstruction used to run the Lagrangian simulations), here it was followed the approach proposed by [11], who set the time window of possible *P. graminis* spore release between 9:00 and 15:00. All the locations crossed by a trajectory out of the above defined time interval were discarded.
6. **Land cover** - Moreover, since the gridded domain considers only those cells covered with a significant wheat surface, also a land cover criterion was implicitly adopted. All those locations whose wheat surface was below the previously defined threshold (1%) were considered as unsuitable.

As a consequence of this selection (Tab. I), from each of the 67.802.400 backward trajectories a set of isolated suitable spore release points (locations) is extracted to define possible connections (Fig. 2). Consecutive suitable release points are linearly interpolated not to omit suitable release locations between two calculated points. For simplicity and for continuity reasons, it was assumed that the segment linking two consecutive suitable locations is, in turn, made of suitable locations.

**TABLE I.**
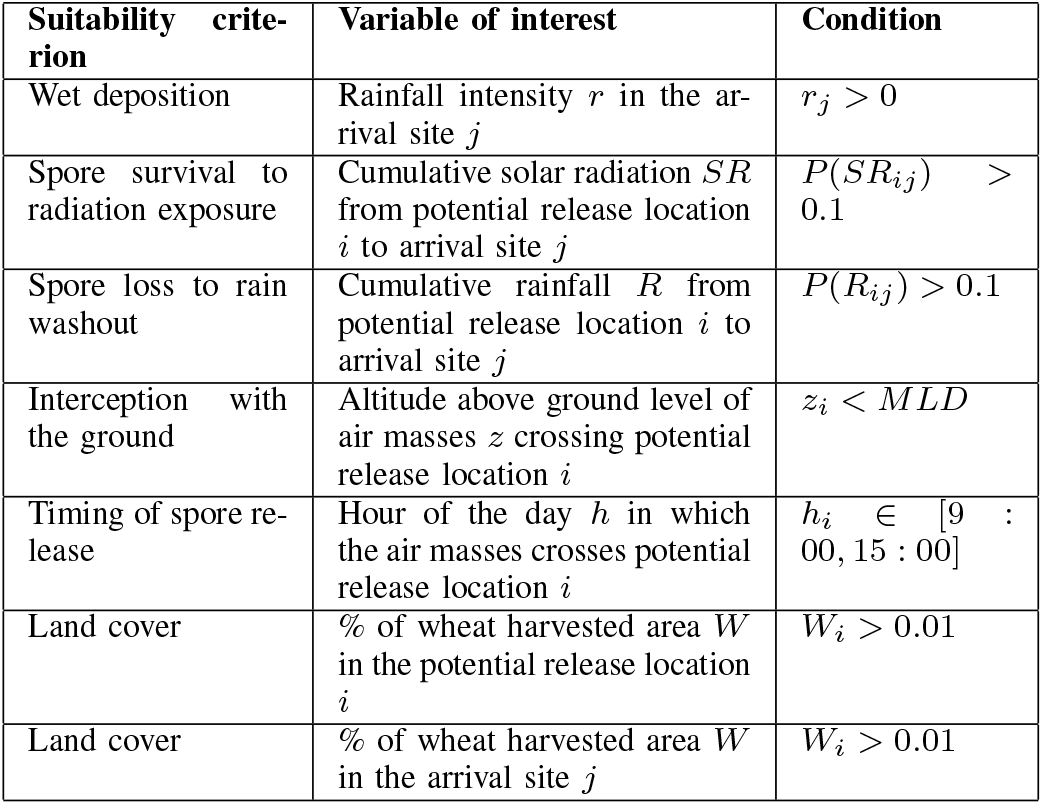
Filtering criteria to identify suitable trajectories

**Fig. 2.**
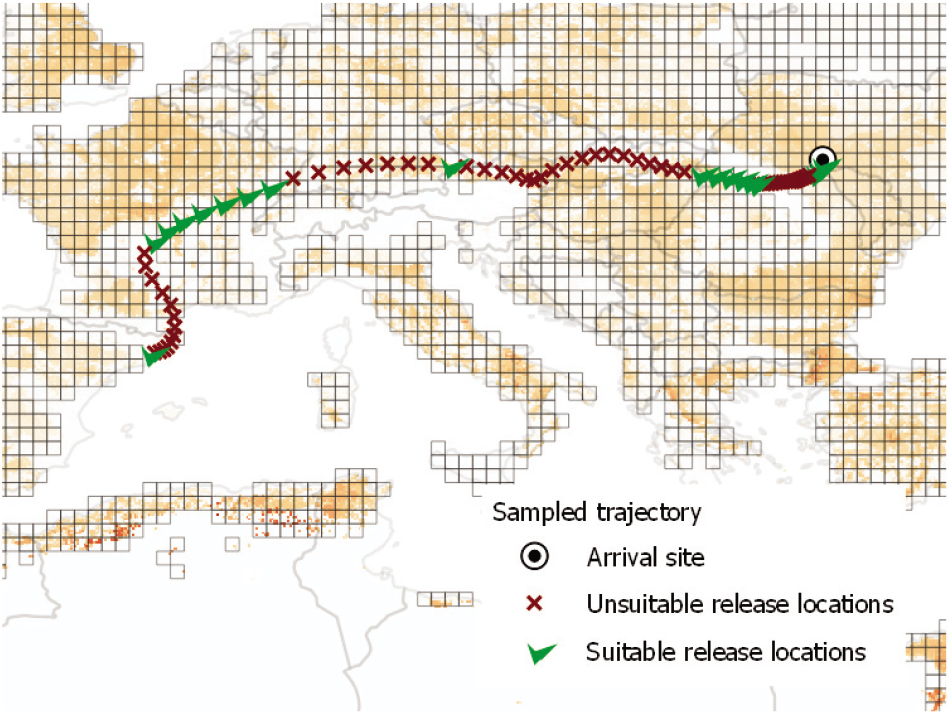
Example of suitable release locations extraction from a backward trajectory whose arrival site is set in western Ukraine. Trajectories are integrated backward from the arrival site. Shades of orange in the map represent wheat density

### D. Daily regional adjacency matrices extraction

After the trajectories were selected, these were intersected to the gridded domain, to obtain 3-hourly boolean adjacency matrices *A* whose elements *a*_*ij*_ show the connections between release locations (rows, *i*) and arrival sites (columns, *j*).

These 17,520 (8 times a day for 6 years) squared 3, 870 × 3, 870 binary contact matrices were firstly grouped and averaged on a week basis, thus obtaining 52 matrices of weekly frequencies of connection, and subsequently aggregated on a geographic basis. Cells have been aggregated according to the second level of the Nomenclature of Territorial Units for Statistics (NUTS1) of the European Union [25]. For simplicity, cells that would be aggregated to a NUTS1 unit with less than two cells have been assigned to the closest NUTS1 region containing two cells or more. Those cells belonging to countries not classified according to the NUTS system were assigned with the name of the country itself, with the exception of Ukraine, Russia and Iran, which have been further divided in more regions because of their size (Fig. 1).

At the and of this spatio-temporal up-scaling, 52 spatial networks were obtained, as weighted and directed adjacency 137 × 137 matrices, whose elements represents the average daily connection frequency for a given week of the year between NUTS1 regions, which will then be considered as the nodes of the geographic networks.

### E. Seasons definition

Temporal structure of wind connectivity patterns was investigated to identify seasonal patterns. Similarity between weekly connectivity matrices was computed by means of the Cut distance algorithm [26] that allows to compare weighted and directed networks having the same number of nodes, and has been used in previous works to identify seasonal wind connectivity patterns [12]. Since the algorithm for computing the Cut distance becomes quickly unfeasible as the number of nodes increases, we used a genetic algorithm to find the pairwise Cut distance between each weekly adjacency matrix. Hence, a hierarchical clustering method based on the Cut distance was used to identify seasonality in the wind connectivity.

### F. Network centrality indicators

The role and the properties of each node within the identified seasonal networks was investigated by means of network centrality indicators. Such indicators reflect the relevance of single nodes within a network with respect to specific dynamics. In particular, such dynamics can be related to epidemic processes or information spreading. Nodes were classified according to the following network centrality indicators:

1. Instrength and outstrength, i.e. the sum of the weights of the edges incoming (resp. outgoing) from a node, which can represent a measure of the potentiality to receive (or spread) a pathogen in a network.
2. VoteRank centrality, which selects a subset of decentralized nodes identified as influential spreaders. In [27] it is shown the performances of this indicator to identify spreaders under Susceptible-Infected-Recovered (SIR) and Susceptible-Infected (SI) models.
3. Coreness, obtained via a weighted k-shell decomposition (WKSD), which assigns a ranking to each layer of nodes that is representative of the proximity to the core of the network, according to its connectivity patterns [29], [30]. Coreness was proved to correlate with nodes with higher spreading potential.
4. Betwenness, which quantifies the number shortest paths between two other nodes of the network passing for a specific node.
5. Closeness, which measure the average distance of a node to all other nodes of the network, and thus its measure of topology centrality.

While the computation of the first indicators was straight-forward, the betwenness and closeness centrality needed a more careful definition. In fact, both indicators rely on some notion of distance between pairs of nodes *i* and *j*, i.e. the shortest path connecting the nodes. While for unweighted networks these distance can be immediately computed as the minimum number of links needed to connect *i* to *j*, for weighted networks several generalized definitions are available depending on the meaning of the weight [31], [32]. In our case the weight of a link represents the frequency of connection, hence it can be considered as a measure of affinity. This means that the distance between nodes *i* and *j* should be linked to a decreasing function of the weight, so that the shortest path will include the heaviest links (the stronger the connection, the lower the distance). In this case, distances between nodes were computed as the reciprocal of the weight [31]:

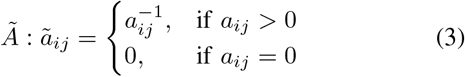

### G. Software

Airmass trajectory extractions were performed using the HYSPLIT software [7], [8] installed on a local cluster (URL cluster), while the rest of computations have been performed with the R software, in particular using the packages sf [33], igraph [34] and GA [35].

## III. Results

### A. Seasonal matrices

The hierarchical clustering method applied to the Cut distance matrix between the 52 weekly networks revealed a clear temporal structure of the connectivity pattern. This structure corresponds to two main seasons: the first one spans for approximately 7 months (from week 14 to week 42, from April to September, hereafter referred as “summer”) and the second covers the remaining 5 months (before week 14 and after week 42, from October to March, hereafter referred as “winter”). These two seasons were hence gathered into two projected static matrices, as the mean of all weekly matrices included in each season.

Since these seasonal networks are characterised by a high edge density (0.606 for summer and 0.779 for winter, computed as the proportion of non-null edges out of all possible edges between the 137 nodes), a filtering method was applied to improve the graphical rendering of the networks. A “backbone” extraction algorithm [36] for the extraction of the statistically relevant connections in weighted and directed network was applied. This method allows to preserve the relevant edges while maintaining the multi-scale nature of local connections. In this case, we run the algorithm trying to preserve at least 20% of the total edge weight. The number of remaining edges after filtering for the summer and winter “backboned” networks is 2.1% (269 out of 12, 97) and 2.9% (428 out of 14, 525) respectively.

The backboned seasonal matrices (Fig. 3 and 4) reveal strong connections in the central-western Europe, with a prevalent longitudinal direction. In the winter backboned matrix, both eastward and westward fluxes are present, but the former appear to be more intense. Eastward connections in central Europe divides in three fluxes; a first, stronger, connects with the Balkan peninsula, were the main axis becomes latitudinal (both northward and southward); a secondary flux deviates northwards towards the Baltic states and Finland, while the last connects to Russia. British islands form a semi-isolated cluster, since they only receive edges from the north of France. They can be seen as a sink with respect of the continental connections. On the opposite, Morocco, Algeria and Tunisia are outward connected to the southern European coasts, while they have no any entering edge. As a consequence, they can be considered a potential source with respect to the continental nodes. Middle East appears to be fully connected, without any preferential connection.

**Fig. 3.**
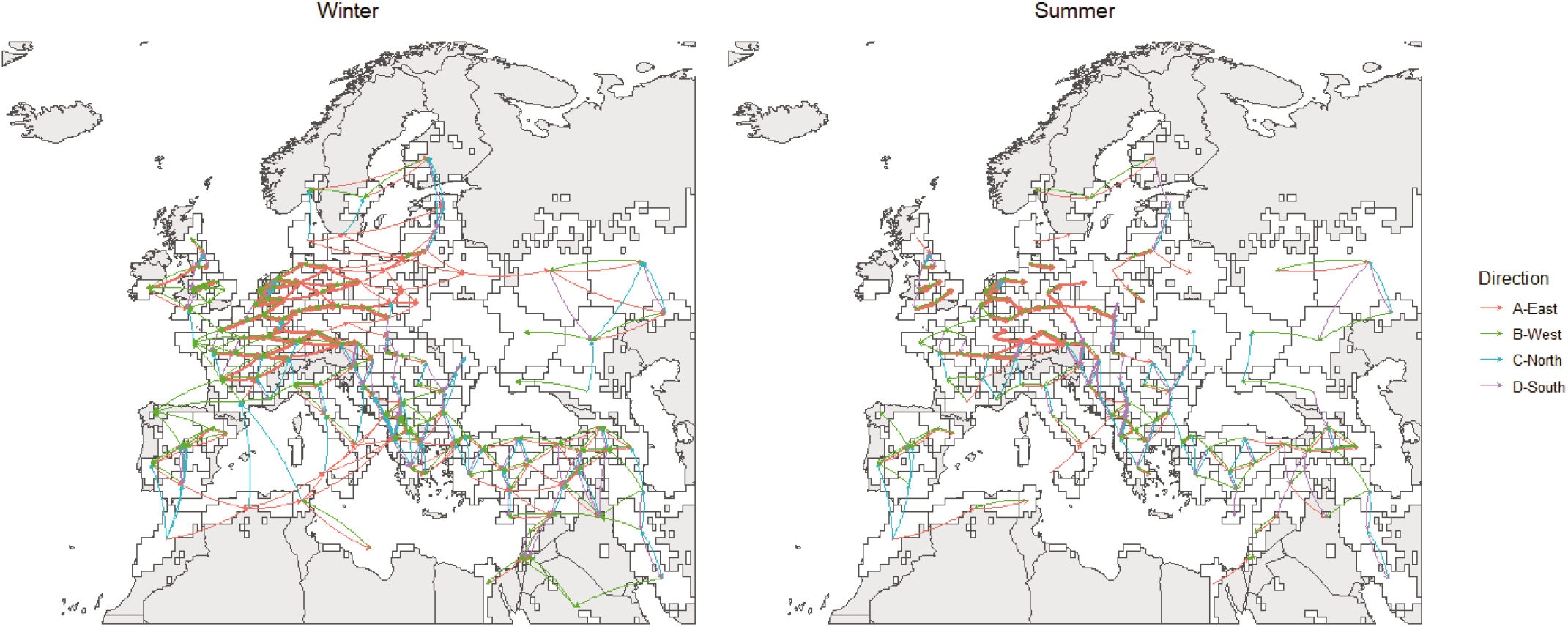
Geographical representation of summer and winter connectivity matrices, as obtained after applying the backbone extraction algorithm [36]. Width of the arrows represents qualitatively the strength of connection, while colours represent the main direction.

**Fig. 4.**
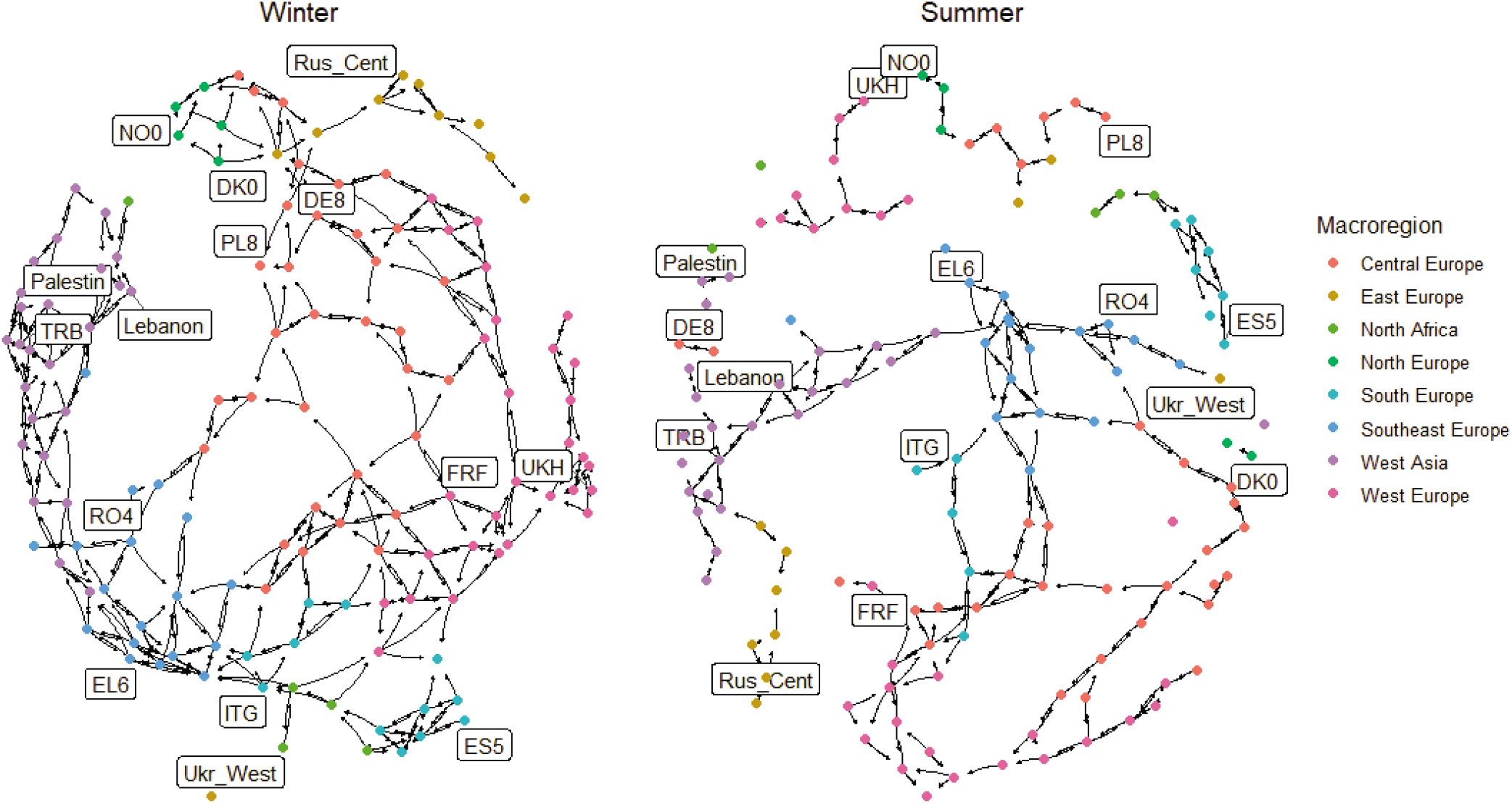
Topological unweighted representation of summer and winter connectivity matrices, as obtained after applying the backbone extraction algorithm [36]. Colours represent the macroregions. A selection of NUTS1 is labelled (cfr. Fig. 1).

The summer network, even though covering a larger period, has a lower density both in the original adjacency matrix and in the backboned one. As a consequence, it is much poorer with connections. Most of them are still shown in central Europe, with a main eastward flux which becomes rapidly southward when approaching the Balkan peninsula. Baltic and Scandinavian countries create an isolated cluster, as well as eastern European countries. British islands are isolated from the continent. Morocco, Algeria and Tunis act again as source towards Europe, but their influence is limited to the Iberian peninsula, which is no longer connected with the rest of Europe. Connections among Middle east countries decrease (for example, Egypt is no longer a sink but only a source, and Saudi Arabia disconnects from other countries).

### B. Network centrality indicators

Network centrality indicators were calculated for all the 137 nodes of the two original seasonal networks (Fig. 5).

**Fig. 5.**
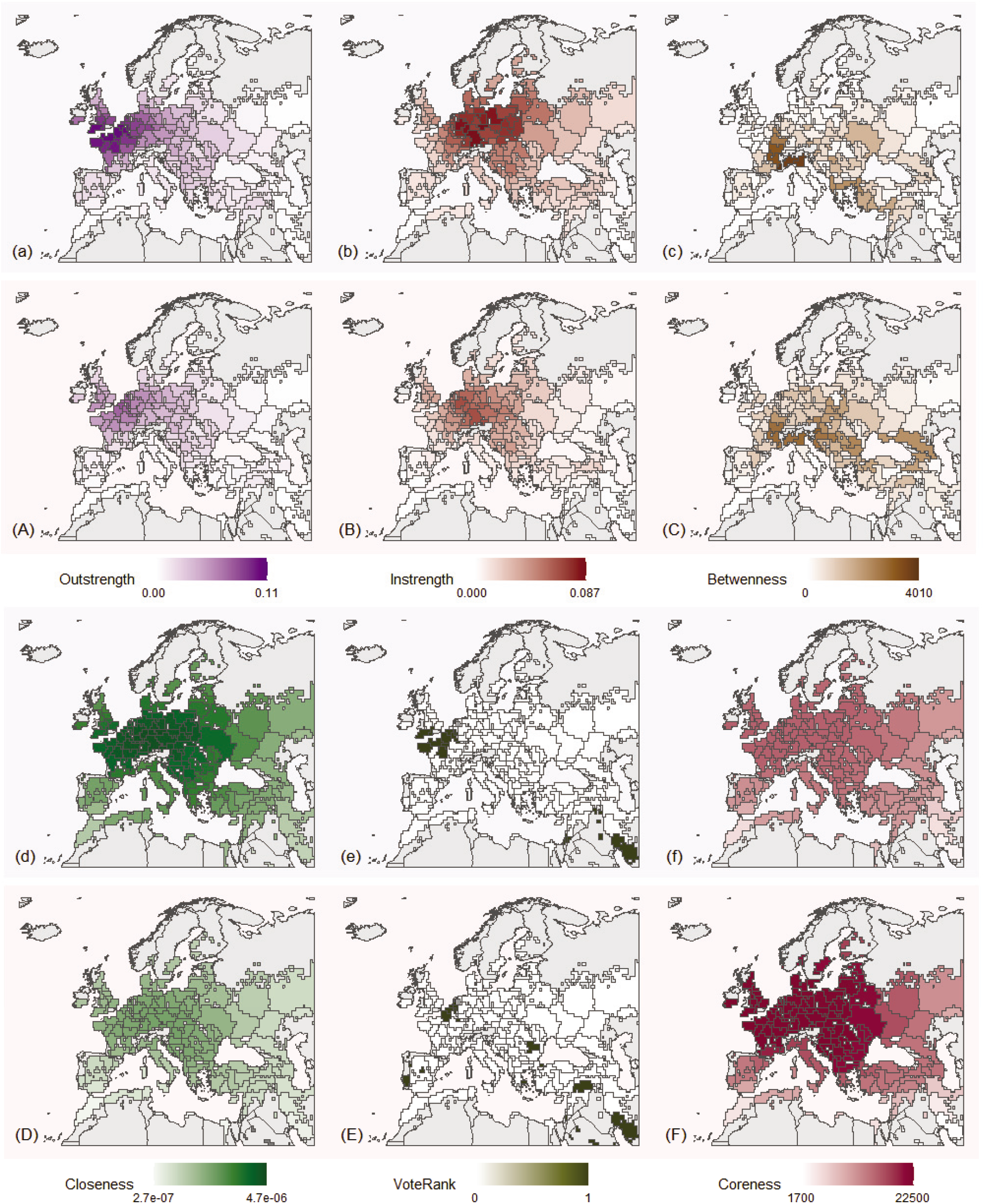
A selection of network centrality indicators: (a) outstrength, (b) instrength, (c) betwenness, (d) closeness, (e) VoteRank, (f) coreness. Blue background (odd rows - lower case letters) is associated with the winter network, red background (even rows - upper case letters) is associated with the summer network.

Outstrength (Fig. 5A) reaches a maximum value of 0.11 during the winter season. In winter, a cluster of nodes with high outstrength on both sides of the English Channel is easily recognisable. In summer, regions characterised by high outstrength are located slightly eastward. In general, regions in central and western Europe are those characterised by the highest values of this indicator.

Instrength indicator (Fig. 5B) reaches a maximum value of 0.87, during the winter season. Nodes characterised with highest instrength are located in Germany and, in general, in central Europe. This pattern appears to be more defined in winter than in summer.

Betwenness (Fig. 5C) identifies different cells compared to outstrength or instrength, highlighting also regions in the geographical periphery. In winter, regions characterised by highest betwenness are located in northern Italy and in south-east France, reaching a maximum value of 4, 010, followed by other regions on the Aegean Sea (Greece, Turkey). In summer, the gradient of betwenness is smoother: regions in northern Italy and in southeast France are confirmed to act as bridges, but also a cluster of regions with medium-high betwenness emerges in the Balkan peninsula and around the coasts of Black Sea (Ukraine, Georgia).

In winter, closeness centrality (Fig. 5D) is higher in western and central European regions. This indicator shows a smoother gradient in the summer compared to winter.

VoteRank (Fig. 5E) algorithm identifies the top *n* nodes which are the best spreaders. In this case, *n* was set to 14, in order to identify the top 10% of the total number of nodes in the network. In winter, there are two definite group of regions identified as spreaders by the VoteRank algorithm. The first is located around the English channel, in Belgium, southern England and, above all, northern France. By contrast, the second is located in the Middle East, and corresponds to Israel, Palestine, Jordan, Iraq and southern Iran. More isolated countries are identified in the summer matrix. There are a group of countries in the Iberian peninsula, another roughly corresponding to Benelux, a third more dispersed in the Middle East (Palestine, Turkey, Saudi Arabia and Iran) and a fourth in the Balkan Peninsula (Romania, Greece, and Istanbul region in Turkey).

Coreness centrality (Fig. 5F), computed via the Weighted K-Shell Decomposition (WKSD) method, identifies layers of regions equally distant from the core. This core-periphery profile is smoother in winter compared to summer. Highest coreness is generally identified in western, central and south-eastern Europe.

### C. Identification of the most influential spreaders based on network properties

Because of the different rankings reflecting different local properties related to network centrality indicators, an analysis aggregating all these indicators was carried out (Fig. 6). In this aggregate analysis, given the different orders of magnitude linked to different meanings of centralities, nodes were characterised in terms of ranking with respect to each indicators rather than absolute values.

**Fig. 6.**
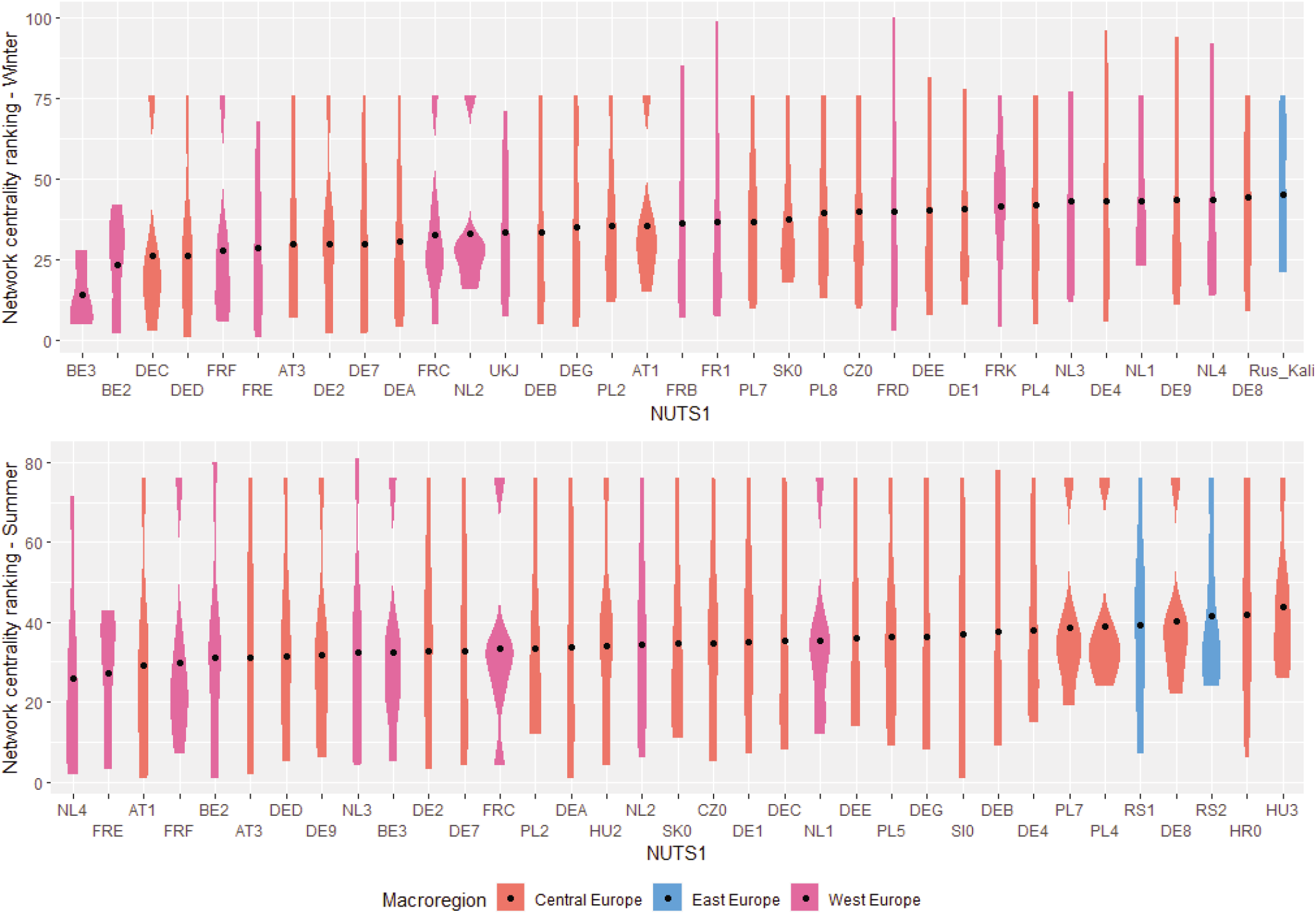
Top 35 NUTS1 regions ranked according to the aggregated computed network indicators. Points represents the average of the rankings for each region. First row: winter. Second row: summer.

Nodes were ordered according to the average ranking in all centrality indicators. Both in summer and in winter, central Europe regions dominates the best performing nodes (20 out of 35 and 24 out of 35, respectively), followed by continental west European regions (14 and 9 out of 35, respectively). Germany and Austria are the most represented country at the top of the rankings among central European regions. However, the very top of the ranking is occupied by west European countries, Flemish and Walloon regions in Belgium in winter and South Netherlands and Hauts-de-France (France) in summer. The only regions outside central and western Europe appearing in this selection are Kaliningrad *oblast’* (Russian exclave *de facto* located in central Europe) in winter and Serbian regions in summer.

## IV. Discussion

Understanding the main features of plant pathogen dispersal is a key step to establish suitable and effective prevention measures in the agricultural context. In this work, features of the spatio-temporal wind connectivity patterns related to potential dispersal of the fungal pathogen *P. graminis* in the European and Mediterranean region were investigated. To perform such analysis, we run several backwards Lagrangian simulations, informed with environmental and meteorological data, to identify potential release and deposition locations and calculate connections frequencies between regions.

Two main seasonal wind connecntion patterns, here referred as winter and summer, were identified. An analysis of both season was carried out, even though wheat rust by *P. graminis* is likely to occur mainly in the first months of the summer season, following the “green wave” of host susceptibility [19]. In both seasons, western and central Europe regions are characterised by strong connections, preferably directed eastwards. Other regions revealed different patterns: northern African countries likely act as a source for southern European countries, while the British islands act more as sink of continental connections; the Balkans (where the main direction of connections is North-South) and the Middle East are generally well connected in multiple directions, while eastern European countries are poorer with connections (Fig. 3). These considerations are more valid in winter, showing denser connectivity, rather than in summer, whose backboned extraction revealed isolated clusters (Fig. 4).

The network analysis stressed the importance of central and western European regions as influential spreaders (Fig. 5 and 6). Regions in central and western Europe have generally high outstrength, instrength, closeness, and coreness. To some extent, VoteRank algorithm, which identifies both central than peripheral nodes, highlights the importance of regions in between of western and central Europe, both in winter and in summer. Betwenness, however, identifies different regions, in particular in the south and south-east of Europe, that could act as “stepping stones” for the propagation front of the disease from the Middle East and East Europe.

Patterns for aerial dispersal of *P. graminis* have been studied or assumed in different parts of the world. *P. graminis* is an obligate pathogen, since its sexual phase requires a alternate host, common barberry (*Berberis vulgaris*) to overwinter [19]. Since barberry have been eradicated in many western countries, it is assumed that spores causing wheat stem rust can enter temperate zones (such as Western Europe or USA) via wind dissemination, after hundreds or thousands of kilometres travels from their regions of origin, within milder continental climates [37]. As a result, in the last century two main infection streams were identified, namely the West European Tract (which origins in Morocco Iberic peninsula and connects up to British islands and Scandinavian peninsula) and East European Tract (which origins in the Lower Danube Plain and drives northward; [38]). A similar stream, named *Puccinia pathway*, was documented in the USA from overwintering southern regions towards northern production area during summer [19]. In recent years, a new highly-virulent variety of stem rust was observed first in Turkey (2005) and developed in a serious outbreak in Germany (2013) from which spread with sporadic infections in Denmark, Sweden and UK [15]. More lately, a large outbreak of stem rust was observed in 2016 in Sicily [15].

The results shown in this work can be read in accordance with the latest observations of stem rust of wheat in Europe. The network analysis revealed the role of Morocco, Algeria and Tunisia as putative sources, and the role of British islands as a sink, confirming the West European Tract. Moreover, the observation of stem rust in Sicily in 2016 suggests even a more important role of north western African countries as a potential source, helped by “stepping stones” in the middle of the Mediterranean. Eventually, the local infections following stem rust outbreak in Germany in surrounding countries confirms the role of central Europe as influential spreader, and the effectiveness of network analysis as a valuable tool to interpret wind borne pathogen dispersal mechanisms.

In this work, Lagrangian simulations coupled with environmental data were used to drive network analysis for the identification of regions potential spreaders of stem rust of wheat. Possible future directions include the refinement of the geographic scale, which is critical to catch local weather condition, the inclusion of wind gusts and temperature at canopy level to indicate conditions for spores release. Moreover, this analysis assumes susceptibility conditions of the host in all the domain, without taking into account agricultural practices or seasonal variability. These could be furthermore integrated with the connectivity networks in a more refined metapopulation SIR model [39], [40].

## V. Acknowledgements

The authors acknowledge the support of funding from the French National Research Agency (ANR) for the BEYOND project (contract # 20-PCPA-0002) and SuMCrop Sustainable Management of Crop Health Program of INRAE that supported the work of all authors, as well as the technical support of Loïc Houde for the computation of HYSPLIT trajectories.

## Notes

https://youtu.be/JIylFAxiu28

### Competing Interest Statement

The authors have declared no competing interest.

